# *Maribacter halichondris* sp. nov., isolated from the marine sponge *Halichondria panicea*

**DOI:** 10.1101/2023.02.14.528435

**Authors:** Leon X. Steiner, Jutta Wiese, Tanja Rahn, Erik Borchert, Beate M. Slaby, Ute Hentschel

## Abstract

A new member of the family *Flavobacteriaceae* (termed Hal144^T^) was isolated from the marine breadcrumb sponge *Halichondria panicea*. Sponge material was collected in 2018 at Schilksee which is located in the Kiel Fjord (Baltic Sea, Germany). Phylogenetic analysis of the full-length Hal144^T^ 16S rRNA gene sequence revealed similarities from 94.3% - 96.6% to the nearest type strains of the genus *Maribacter*. The phylogenetic tree depicted a cluster of strain Hal144^T^ with its closest relatives *Maribacter aestuarii* GY20^T^ (96.6%) and *Maribacter thermophilus* HT7-2^T^ (96.3%). Genome comparisons of strain Hal144^T^ with *Maribacter* spp. type strains exhibited average nucleotide identities in the range of 75% - 76% and digital DNA-DNA hybridization values in the range of 13.1% - 13.4%. Strain Hal144^T^ was determined to be Gram-negative, mesophilic, strictly aerobic, flexirubin positive, resistant to aminoglycoside antibiotics, and able to utilize N-acetyl-β-D-glucosamine. Optimal growth occurred at 25 – 30 °C, within a salinity range of 2 - 6% sea salt, and a pH range between 5 - 8. The major fatty acids identified were C_17_:_0_ 3-OH, iso-C_15_:_0_, and iso-C_15:1_ G. The DNA G+C content of strain Hal144^T^ was 41.4 mol%. Based on the polyphasic approach, strain Hal144^T^ represents a novel species of the genus *Maribacter*, and we propose the name *Maribacter halichondris* sp. nov.. The type strain is Hal144^T^ (= DSM 114563^T^ = LMG 32744^T^).

## Introduction

The genus *Maribacter* (*Bacteroidota, Flavobacteriia, Flavobacteriales, Flavobacteriaceae*) comprised 29 validly described species and two not yet validated species at the time of writing [1]. Most species were derived from marine sources such as sponges [2], red and green algae [3,4], seawater [5], sediments [6], and tidal flat [7]. Few metabolic features indicating adaptation of members of the genus *Maribacter* to environmental conditions were described. Among them are the production of carbohydrate-active enzymes, such as agarase, alginate lyase, carrageenase, glycoside hydrolases, pectate lyase, polysaccharide lyases, and xylanase being important for habitats, where phytoplankton and macroalgae produces diverse polysaccharides [8,9,10]. Tolerance to heavy-metals, such as Co^2+^ (10mM) and Cd^2+^ (0.5 mM) was reported for *Maribacter cobaltidurans* B1^T^, which was isolated from deep-sea sediment [11]. Only very little is known about the biological role of *Maribacter* spp. Strains in host-microbe interactions. *Maribacter* sp. MS6 drives symbiotic interactions with the green macroalga *Ulva mutablis* by releasing morphogenetic compounds, e.g. the hormone like compound thallusin, which aid in algal morphogenesis, such as rhizoid and cell-wall formation [12,13]. A *Maribacter* sp. strain reduces the reproductive success in the diatom *Seminavis robusta* [14]. Recently, *Maribacter* sp. strains closely related to *Maribacter dokdonensis* DSW-8^T^ and *Maribacter sedimenticola* KMM 3903^T^ with > 98.50 % similarity of 16S rRNA gene sequences were isolated from sponges collected from the Pacific Ocean [15]. These *Maribacter* spp. isolates showed antibiotic activity against *Mycobacterium smegmatis*. Three Maribacter isolates from the sponge *Hymeniacidon perlevis* sampled at Nord-Pas-de Calais (France) showed antibacterial effects against multi-drug resistant *Staphylococcus aureus*. These isolates were affiliated to *Maribacter arcticus* KOPRI 20941^T^ with approx. 98.50 % similarity of 16S rRNA gene sequences [16]. We are currently developing the Baltic Sea sponge *Halichondria panicea* as an experimental model for marine sponge-microbe-phage interactions [17], Steiner *et al*. unpublished]. *H. panicea* inhabits coastal areas around the globe and harbors a diverse microbial community including the symbiont *Candidatus* Halichondribacter symbioticus [18,19]. Our bacterial cultivation approaches from this sponge species resulted in more than 350 isolates, including 7 *Maribacter* spp. strains. Among them, strain Hal144^T^ attracted our attention as it represents a putatively novel species and serves as a host strain for a novel phage (Steiner *et al*. unpublished). The present study identifies the taxonomic status of strain Hal144^T^ by determining its phylogenetic, physiological, and genomic properties.

### Isolation and ecology

Strain Hal144^T^ was isolated as part of a larger microbial community analyses from the marine breadcrumb sponge *Halichondria panicea*. Sponge individuals were sampled via snorkeling on October 2^nd^, 2018 from Kiel, Schilksee (Baltic Sea, Germany, coordinates: latitude 54.424705, longitude 10.175133). Specimens were transported in 500 ml Kautex bottles to GEOMAR Helmholtz Centre for Ocean Research Kiel within 2 hours after collection. 8.8 g sponge material was rinsed three times with 0.2 µm filtrated and autoclaved Baltic Sea water (BSW) to remove loosely attached particles and microorganisms. The sponge sample was homogenized in a 50 ml Falcon plastic tube, with 35 ml BSW by use of an Ultraturrax for 30 seconds at 17,500 rpm and serially diluted with BSW from 10^−1^ to 10^−4^. 100 µl of the undiluted suspension and of the dilutions were spread onto a tryptone containing medium (1 g tryptone, 1 g yeast extract, 15 g Bacto-Agar, 1000 ml Baltic Sea water, pH 7.5) and incubated at 25°C for 7 days. Hal144^T^ was obtained from a colony growing on the dilution 10^−4^ and cultured on tryptone agar plates at 25°C, then subcultivated using marine medium (MB, 37.4 g BD Difco™ Marine Broth 2216 (Becton Dickinson and Company, New Jersey, USA), 15 g Bacto-Agar, 1000 ml aq. deion.) for 7 days at 25°C before cryopreservation with the Cryobank System (Mast Diagnostica GmbH, Reinfeld, Germany) at -20 °C and -80°C. This study aimed to taxonomically describe Hal144^T^ using morphological, physiological, phylogenetic, and genomic characteristics.

### 16S RNA phylogeny

Genomic DNA of strain Hal144^T^ was extracted using the DNeasy Blood & Tissue Kit (Qiagen GmbH, Hilden, Germany) according to the manufacturer’s instructions. The 16S rRNA gene sequence was amplified using the primers Eub27F (5’-GAG TTT GAT CCT GGC TCA G-3’) [20] and Univ1492R (5’-GGT TAC CTT GTT ACG ACT T-3’) [21] sequenced via Sanger sequencing [22] at Eurofins Genomics (Ebersberg, Germany) with the primers 534R [23], 342F [24], and Univ1492R [21]. The sequenced contigs were assembled and the quality of the sequence was assessed using ChromasPro 2.1.8 (Technelysium Pty Ltd, Brisbane, Australia). The partial 16S rRNA gene sequence comprised 1488 base paires (bp) and was deposited under the accession number MT406525.2. This PCR-based 16S rRNA gene sequence is identical with full-length genome-derived sequence (1531 bp) in the overlapping region. The full-length 16S rRNA gene sequence of Hal144^T^ was aligned to all *Maribacter* spp. type strains and *Capnocytophaga ochracea* DSM 7271^T^ as outgroup using the ClustalW tool of MEGA version 11.0.13 [25]. Phylogenetic trees were constructed using the Neighbor-Joining (NJ) method [26] and computing the evolutionary distances with the Maximum Composite Likelihood method [27], the minimum evolution (ME) method in combination with the Maximum Composite Likelihood method and Close-Neighbor-Interchange (CNI) algorithm [27,28,29], and the Maximum-Likelihood (ML) method in combination with the Tamura-Nei model [30], to ensure the consistency of the tree topology. The phylogenetic trees were constructed by MEGA 11.0.13 [25] by running 1000 bootstrap replications and including 1^st^+2^nd^+3^rd^+noncoding positions [31]. The resulting trees were drawn to scale, with branch lengths measured in the units of the number of base substitutions per site.

Phylogenetic 16S rRNA gene sequence analysis revealed that the strain Hal144^T^ affiliated to the genus *Maribacter*). Applying the NJ (Fig. 1), ME (data not shown), and ML (data not shown) method, strain Hal144^T^ clustered with *Maribacter aestuarii* GY20^T^ (= JCM 18631^T^) (96.57%). This cluster branched with *Maribacter thermophilus* HT7-2^T^ (96.31%). 16S rRNA gene sequence similarity to all *Maribacter* spp. type strains is in the range from 94.26% to 96.57% indicating that strain Hal144^T^ belongs to a new species according to a < 98.7% threshold [33].

**Figure 1.**
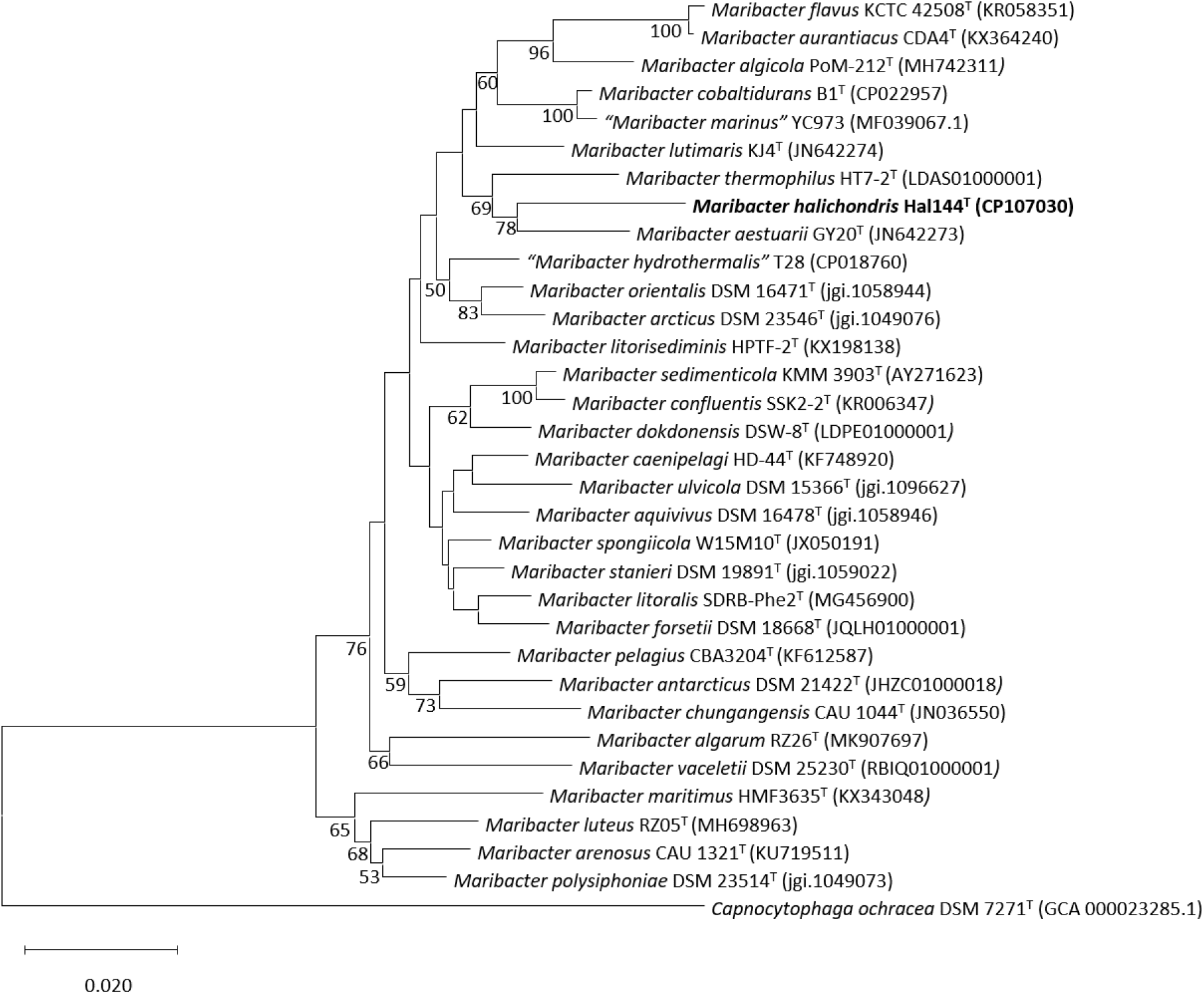
Phylogenetic relationships of Hal144^T^ based on 16S rRNA gene sequences using the Neighbor-Joining method. As far as available, whole 16S rRNA gene sequences were derived from genome sequences applying ContEst16S [32]. Bootstrap values (≥50 %) based on 1000 replications are shown next to the branches. A total of 1540 positions were in the final dataset. *Capnocytophaga ochracea* DSM 7271^T^ was used as outgroup. Bar, 0.02 substitutions per nucleotide position.

### Genome features

Strain Hal144^T^ and *M. aestuarii* JCM 18631^T^ (= GY20^T^), (obtained from RIKEN BioResource Research Center, Tsukuba, Japan), were grown on MB at 25 °C for 7 days. DNA was extracted with Qiagen Genomic-tip 100/G (Hilden, Germany), following the standard protocol by the manufacture. The extracted DNA had a concentration of 279 ng/µl in case of Hal144^T^ and 397 ng/µl for JCM 18631^T^. The quality of the DNA met the criteria, i.e. A260/280 ratio of >1.8 and A260/230 ratio of <1.8, according to NanoDrop (Thermo Fisher Scientific, Germany) measurements. The genome was sequenced with MinION nanopore technology (Oxford Nanopore Technologies, Oxford, UK) using a MinION Flongle Flow-Cell (Cat.No. FLO-FLG001) with the Flow Cell Priming Kit (Cat.No. EXP-FLP002) and the Rapid Sequencing Kit (Cat.No. SQK-RAD004), following the protocols by the manufacturer. The super-accurate model of Guppy (Oxford Nanopore Technologies plc. Version 6.2.1+6588110, dna_r9.4.1_450bps_sup) was used for basecalling of the MinION reads. Initially, the MinION data were assembled using Miniasm (version 0.3-r179) [34], then polished with Racon (version 1.5.0) [35] and Medaka (version 1.4.3, model r941_min_sup_g507) [36]. The annotation was prepared using RAST [37]. The GenBank accession numbers for the genome sequences of strain Hal144^T^ and of *Maribacter aestuarii* JCM 18631^T^ are CP107030 and CP107031, respectively.

The general genomic features were determined using Quast 5.2 [38], Prokka 1.3 [39], and CheckM [40] and are displayed in Table 1. DNA G+C content was 41.4 mol% (Table 1), which is in the range of 35 % - 41.8 % as it was calculated for all 29 *Maribacter* spp. type strains with Quast in this study. The range 35 % - 39 % given in the description of the genus *Maribacter* [42] based only on 8 *Maribacter* species. The average nucleotide identities (ANI) were determined using the ANI calculator from the enveomics collection [43]. ANI values between strain Hal144^T^ and *Maribacter* spp. type strains were in the range of 75 % - 76 %. Since these mean identities are below the threshold (95 - 96 %) for species delineation [44,45], strain Hal144^T^ represents a novel species.

**Table 1.**
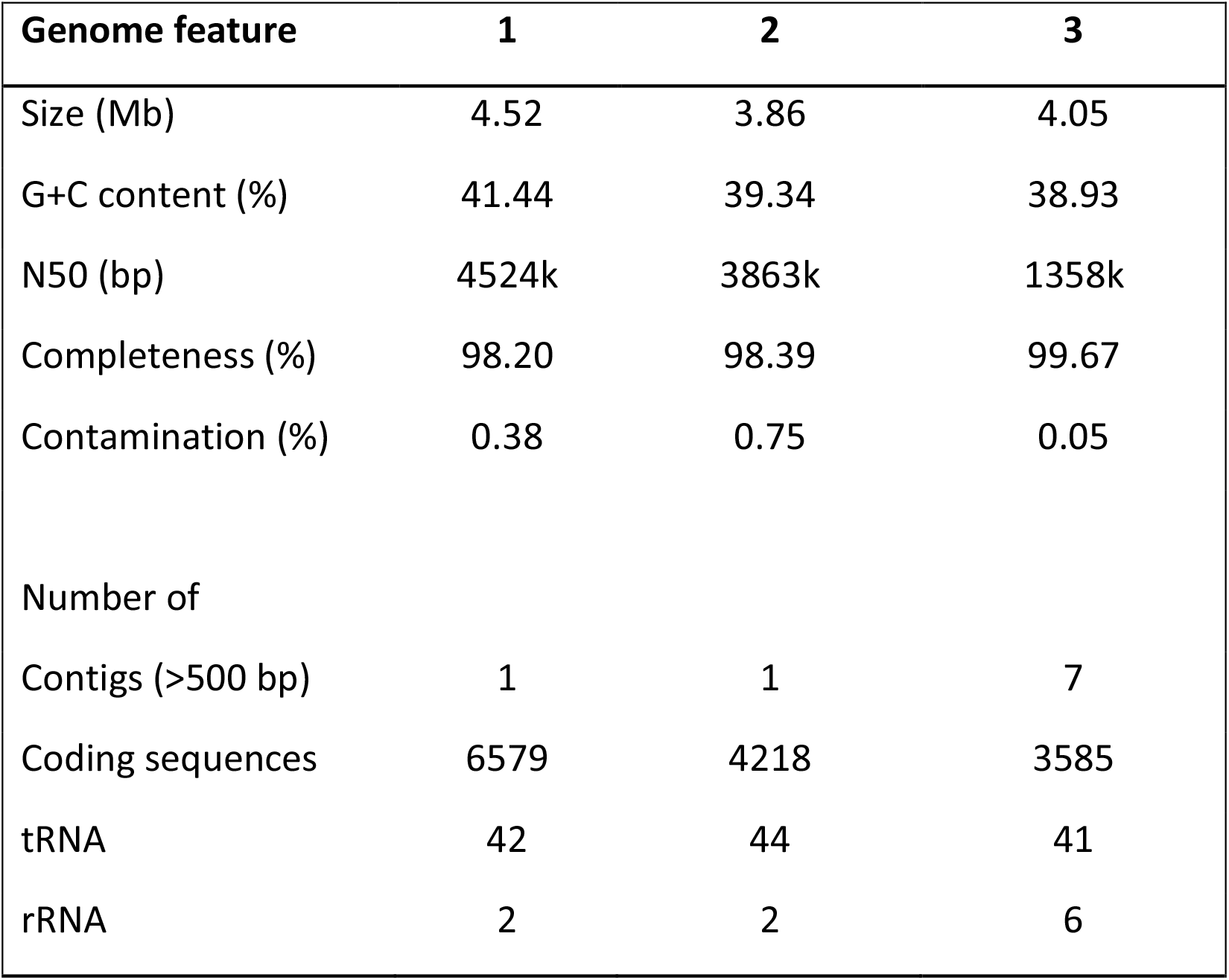
Comparison of the general genomic features of strain Hal144^T^ and related species of the genus *Maribacter*. Strains: 1, Hal144^T^ (CP107030, this study); 2, *Maribacter aestuarii* JCM 18631^T^ (CP107031, this study); 3, *Maribacter thermophilus* HT7-2^T^ (GCF_001020565), [41]).

Digital DNA-DNA hybridization (dDDH) values were calculated using the dDDH calculator provided on the platform of the Type (Strain) Genome Server (TYGS) [46]. The mean dDDH values determined for strain Hal144^T^ compared to type strains of the genus *Maribacter* were in the range of 13 % - 13.4 %, all below the suggested boundary (< 70%) for species delineation [47], demonstrating that strain Hal144^T^ represents a novel genomic species.

Genome-based phylogeny was calculated with strain Hal144^T^ and with publically available *Maribacter* spp. genomes. Two *Maribacter* spp. clusters branched from the outgroup (Fig. 2). Applying the NJ-method (Fig. 2), the ME-method (data not shown), and the ML-method (data not shown) one cluster contained *Maribacter algarum* RZ26^T^, an isolate from the red alga *Gelidium amansii, Maribacter vaceletii* W13M1A^T^, an isolate from the sponge *Suberites carnosus*, and also strain Hal144^T^. This cluster is separated from the clade with *Maribacter luteus, Maribacter arenosus*, and *Maribacter polysiphoniae*. The differences in the 16S rRNA gene sequence phylogeny and the genome-based phylogenetic trees might be a result of the various target proteins used for the calculations, i.e. one 16S rRNA gene versus 120 single copy marker genes. Further, the different numbers of available sequences for the type strains, i.e. 25 genome sequences versus 32 16S rRNA gene sequences, may have led to divergent phylogenies. It is expected, that the comparison of phylogenetic trees will become more meaningful, when more genomic data are available for diverse *Maribacter* spp. type strains.

**Figure 2.**
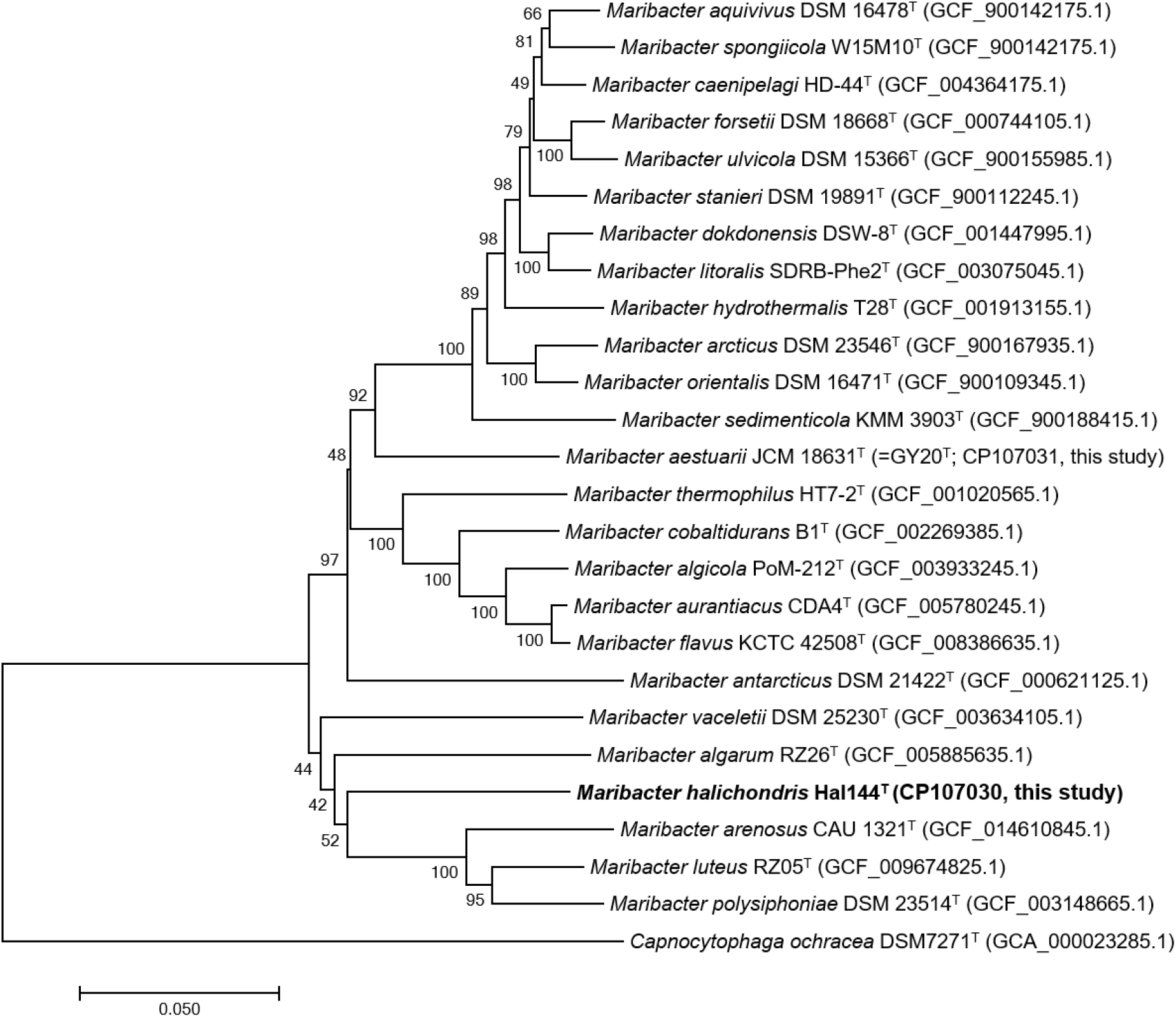
Genome phylogeny of strain Hal144^T^ was inferred using the GTDBtk pipeline [48,49]. The pipeline was used with its version 2.1.0 and is based on 317,542 reference genomes. For bacterial genomes, the taxonomic identification is based on 120 single copy marker proteins. The pipeline employs MEGA version 11.0.13 [Tamura et al. 2021] to calculate the phylogenetic trees using the Neighbor Joining method. Bar, 0.05 substitutions per nucleotide position.

The Gram-negative strain Hal144^T^ exhibited resistance against the antibiotics ampicillin, bacitracin, mupirocin, and oleandomycin, which are mainly used against Gram-positive bacterial infections. Resistance to antibiotics against Gram-negative bacteria was observed. Among them were all five aminoglycosides tested in this study i.e. amikacin, kanamycin, gentamicin, neomycin, and streptomycin, and the polypeptide polymyxin B. Membrane-associated antibiotic resistance, e.g. by the expression of multidrug efflux pumps, is a key mechanism in Gram-negative bacteria [50,51]. Therefore, genome analysis focused on related characteristics. Applying the RAST (Rapid Annotations using Subsystems Technology) - Server [37], two multidrug resistance efflux pumps families were detected. One efflux pump system belongs to multi antimicrobial extrusion protein (MATE), a Na^+^/drug antiporter. The second system, called resistance nodulation division (RND), a H^+^/drug antiporter, confers resistance to antimicrobial compounds produced by the host and plays a role in colonization, persistence, and dissemination of bacteria in the host [51]. In addition, a gene coding for CmeC, an outer membrane channel protein originally described in efflux systems from *Campylobacter* spp. strains [50], was found.

Chitin, a polymer of N-acetyl-β-D-glucosamine (GlcNAc), is the most abundant biopolymer in the marine environment [52] and it is an important structural component within the structural fibers of sponges belonging to the class Demospongia [53]. It is assumed, that *H. panicea* also contains chitin, which could potentially be cleaved by sponge-associated bacteria with endo- and exo-chitinases into oligomers and dimers [54]. Hal144^T^ is able to produce the monomer GlcNAc by its N-acetyl-β-glucosaminidase activity. RAST – server [37] and web-resources of the Bacterial and Viral Bioinformatics Resource Center (BV-BRC) [55] were used to prove the presence of enzymes and transporters involved in the utilization of GlcNAc in strain Hal144^T^. GlcNAc from the environment might be transferred into the periplasm by an outer membrane protein (OmpA). The following four transport systems for GlcNAc from the periplasma to the cytoplasma were identified: (i) N-acetylglucosamine-specific phosphotransferase system (EC 2.7.1.69) consisting of the IIA (NagEa), IIB (NagEb), and IIC (NagEc) component, (ii) N-acetylglucosamine transporter (NagP), (iii) N-acetylglucosamine related transporter (NagX), and (iv) ATP-binding cassette (ABC) N-acetyl-D-glucosamine transporter system consisting of ATP-binding protein (ABCa), permease protein 1 (ABCb1), permease protein 2 (ABCb2), and sugar-binding protein (ABCc). The NagE system releases GlcNAc-6P. Acetate is cleaved by N-acetylglucosamine-6-phosphate deacetylase (NagA, EC 3.5.1.25) and glucosamine-6-phosphate deaminase (NagB1 and NagB2, EC 3.5.99.6) metabolizes glucosamine-6P to fructose-6P by cleaving ammonia. Fructose-6P is further processed in the glycolysis.

### Physiology and chemotaxonomy

The morphological characteristics of strain Hal144^T^ were assessed using 7-day old cultures incubated on MB medium at 25 °C. Colony morphology and color were evaluated via observation with a loop, while cell morphology and motility were examined via light microscopy (Carl Zeiss Axiophot epifluorescence microscope). The results are depicted in the species description. Gram-staining was performed using the bioMérieux Color Gram 2 Test Kit (bioMérieux Deutschland GmbH, Nürtingen, Germany) according to the manufacturer’s instructions and showed strain Hal144^T^ to be Gram-negative.

The physiological and biochemical characteristics were also studied. Salinity-dependent growth was determined with 1 % intervals of both, 0 - 7 % (w/v) NaCl and 0 - 7 % (w/v) Tropic Marine sea salt classic (Wartenberg, Germany), on a medium with the following ingredients: 5.0 g BD Bacto™ Peptone, 1.0 g BD Bacto™ Yeast Extract, 15.0 g BD Bacto™ Agar, 1 L of deionized water. The cultures were incubated at 25 °C for 7 days. Temperature-dependent growth was assessed at 5 °C - 40 °C (intervals of 5 °C) on MB for 7 days. pH-dependent growth of the strains was assessed at 5.0, 6.0, 6.5, 7.5, 8, 8.5, 9.0, and 9.5 on MB at 25 °C for 7 days with the addition of 1 M NaOH and 1 M HCl solutions to adjust the pH level. Specific enzymatic activities were studied using the semi-quantitative API^®^ ZYM test kit (bioMérieux) according to the manufacturer’s instructions using 0.9 % NaCl solution as an inoculum. The test strips were incubated for a period of 18 h at 25 °C in the dark. 3 % (v/v) hydrogen peroxide was added to colonies of the strains and the formation of gas bubbles [56] was observed to determine catalase activity. Oxidase activity was tested by smearing colonies onto a non-impregnated filter paper disc soaked with bioMérieux oxidase reagent (N,N,N,N-tetramethyl-1,4-phenylenediamine) and observing the development of a violet to purple coloration within 10 -30 sec according to the manufacturer’s instructions. Oxygen requirements were assessed with the aerobic/anaerobic test tube method [57] using soft agar MB medium (7.48 g Difco™ Marine Broth 2216, 1.2 g BD Bacto™ Agar in 200 ml of deionized water) and incubation at 25 °C for 1 week. The presence of pigments was also investigated. The KOH test was performed with 7-day old cultures of the strain Hal144^T^ to detect the presence of flexirubin-type pigments [58]. The phenotypic results of Hal144^T^ are displayed in the species description.

Growth of strain Hal144 was assessed at 25°C on MB agar plates in comparison to liquid MB medium (100 ml MB in 300 ml Erlenmeyer flasks with three baffles, 120 rpm). Only with a high amount of inoculum (approximately a half culture agar plate) growth was observed in liquid media. The bacterial cells were not homogeneously distributed in liquid media. Instead, the cells formed crumbs, which attached ring-shaped on the glass wall in the aerated zone. It is assumed, that strain Hal144^T^ prefers surfaces for growth, at least when subjected to the cultivation conditions in our study. This could indicate, that strain Hal144^T^ contributes to the formation of microbial biofilms in its host *Halichondria panicea* and thus might play a role in the complex cellular dialogue of the sponge holobiont [59].

Cellular fatty acids of strain Hal144^T^ were analysed by DSMZ Services (Leibniz Institute DSMZ, Braunschweig, Germany) using a 7-day old culture grown on MB medium at 25 °C. Briefly, fatty acid methyl esters were obtained by saponification, methylation, and extraction using minor modifications of the methods of Miller *et al*. [60] and Kuykendall *et al*. [61]. The fatty acid methyl mixture was separated using a device consisting of an Agilent 7890B gas chromatograph fitted with a 5% phenyl-methyl silicone capillary column (0.2 mm × 25 m), a flame ionization detector, an Agilent model 7683A automatic sampler, and a HP-computer with MIDI data base (Hewlett-Packard Co., Palo Alto, California, U.S.A.). The Sherlock Microbial Identification System (MIS) Standard Software (Microbial ID, MIDI Labs inc, Newark, Delaware, U.S.A) automatical integrated the peaks, identified the fatty acids, and calculated their percentage content using the TSBA6 database. The fatty acid profile is shown in Table 2. The major fatty acids observed were iso-C_17_:_0_ 3-OH, iso-C_15_:_0_, and iso-C_15:1_ G, followed by fatty acids from the category “summed feature 3” consisting of C_16 :1_ω6c and/or C_16 :1_ω7c. The overall fatty acid pattern of strain Hal144^T^ was similar to those described for other *Maribacter* spp. type strains [42].

**Table 2.**
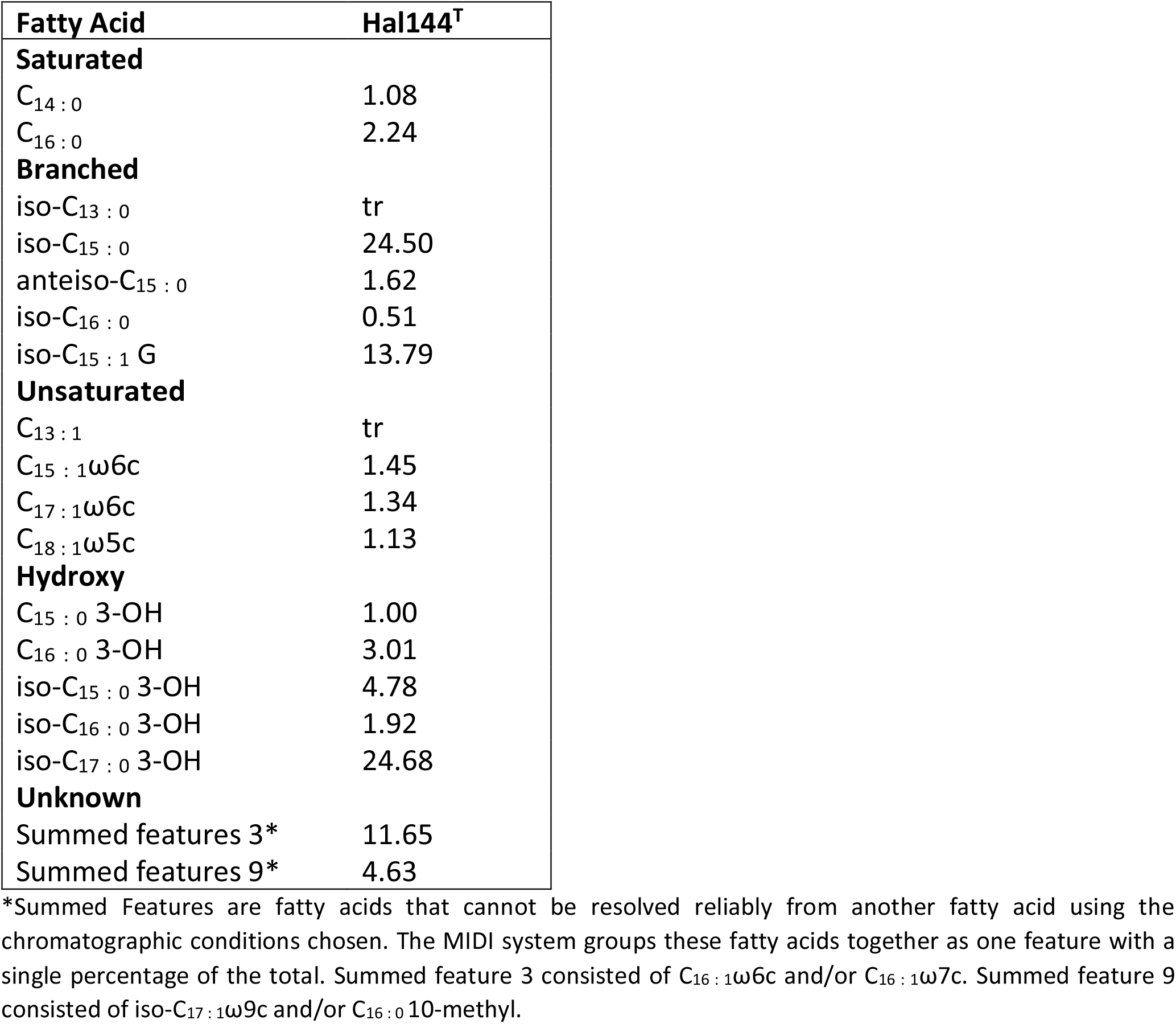
Cellular fatty acid composition (%) of Hal144^T^. tr, trace (less than 0.5 %).

The sensitivity / resistance to 29 antibiotics was tested using the disc diffusion [62]. The test was performed with Oxoid antimicrobial susceptibility test discs (Otto Nordwald GmbH, Hamburg, Germany) on MB medium, which was inoculated with a 7-day culture of strain Hal144^T^ using a swab (bioMérieux). In addition, the effect of trophodietic acid (TDA) on the growth of strain Hal144^T^ was determined, since the compound exhibited antimicrobial activity against clinical pathogens, while TDA resistance was observed in marine bacterial isolates belonging to different taxa, including the genus *Maribacter* [63]. TDA (<u>AdipoGen Life Sciences</u>, Fuellinsdorf, Switzerland) was solved in methanol, dropped on antibiotic test discs (Ø 6 mm, Machery-Nagel, Düren, Germany), and the methanol was evaporated before placing the test disc on the culture. Plates were incubated for 7 days at 25°C. The presence of a clear inhibition zone around the test disc indicated the susceptibility to the tested antibiotic. The antibiotic profile of strain Hal144^T^, including the respective test concentration, is presented in the species description.

### Protologue

#### Description of *Maribacter halichondris* sp. nov

ha.li.chon’dris. N.L. masc. adj. *halichondris*, from Greek χαλις, silica, from Greek χονδρος chondral, pertaining to *Halichondria*.

Cells are Gram-negative, strictly aerobic, non-motile, stabs or slightly curved stabs, 0.8 µm wide and 2 µm long. Colonies are circular, raised, orange, and brittle, 1 - 2 mm in diameter. Growth occurs on 2 - 6 % (w/v) sea salt (optimum 3 - 4 %), no growth on NaCl as the sole salt supplement, at 5 °C - 30 °C (optimum 25 °C - 30 °C), and at pH 5.0 - 8.0 (optimum pH 6.5 - 7.5). Strain is oxidase- and catalase-positive. Positive for alkaline phosphatase, esterase (C4), esterase lipase (C8), lipase (C14), leucine arylamidase, valine arylamidase, cystine arylamidase, α-chymotrypsin, acid phosphatase, naphthol-AS-BI-phosphohydrolase, and N-acetyl-β-glucosaminidase, and weak positive for β-galactosidase, α-glucosidase, β-glucosidase, α-mannosidase, and trypsin, but negative for α-galactosidase, β-glucuronidase, and α-fucosidase. Displays sensitivity to cefoxitin (30 µg), chloramphenicol (50 µg), ciprofloxacin (5 µg), doripenem (10 µg), doxycycline (30 µg), imipenem (10 µg), linezolid (30 µg), norfloxacin (10 µg), novobiocin (30 µg), ofloxacin (5 µg), rifampicin (30 µg), teicoplanin (30 µg), tetracycline (30 µg), and vancomycin (30 µg). Exhibits resistance to amikacin (30 µg), ampicillin (10 µg), bacitracin (10 units), kanamycin (30 µg), mupirocin (200 units), gentamicin (30 µg), neomycin (30 µg), oleandomycin (15 µg), polymyxin B (300 units), streptomycin (25 µg), and trophodietic acid (2 µg). Variable reactions for erythromycin (15 µg), nalidixic acid (30 µg), lincomycin (15 µg), oxacillin (5 µg), and penicillin G (10 units). Flexirubin-type pigment is present. The major fatty acids (> 5 % of total composition) are C_17_:_0_ 3-OH, iso-C_15_:_0_, and iso- C_15:1_ G. The DNA G+C content of Hal144^T^ is 41.4 mol%.

The type strain Hal144^T^ (= DSM 114563^T^ = LMG 32744^T^) was isolated from the marine sponge *Halichondria panicea* collected at Schilksee along the Kiel-Fjord of the Baltic Sea (latitude 54.424705, longitude 10.175133).

## Author statements

### Author contributions

L.X.S.: Conceptualization, method establishment, formal analyses (genomics); J.W.: Conceptualization, validation, writing original draft; E.B.: Formal analyses (assembling, genomics); T.R.: Formal analyses (microbiology); B.M.S.: Method establishment, assembling, validation; U.H.: Conceptualization, supervision, revision; All authors were involved in the processes of writing and reviewing.

### Consent for publication

The authors are consent to publication.

## Acknowledgements

We acknowledge Dr. Lara Schmittmann for sponge collections and for on-going discussions.

## Abbreviations

ABC: ATP-binding cassette
BCCM/LMG: Belgian Coordinated Collections of Microorganisms
BSW: Baltic Sea water
DSMZ: Leibniz-Institut DSMZ-Deutsche Sammlung von Mikroorganismen und Zellkulturen
GlcNAc: N-acetyl-β-D-glucosamine
MB: marine medium
rpm: rounds per minute.

